# *L*-5-[11C]-glutamine PET of Breast Cancer: Kinetic Analysis in Mouse Models to Evaluate Glutamine Metabolism

**DOI:** 10.64898/2026.07.02.736194

**Authors:** Raheema Damani, Christopher Hensley, Hoon Choi, Hsiaoju Lee, Rong Zhou, Austin R. Pantel, David A. Mankoff, Elizabeth J. Li

## Abstract

**Background:** Glutamine addiction is a hallmark of aggressive tumors, yet *glutaminase (GLS1)* inhibitor CB-839 showed disappointing anti-tumor efficacy in clinical trials. *L*-5-[¹¹C]-glutamine ([¹¹C]glutamine) PET enables non-invasive assessment of glutamine metabolism *in vivo*, providing a tool to test mechanistic hypotheses, and identify tumors likely to respond to *GLS1* inhibition: focusing on compartmentation of *GLS1-*derived glutamate, CB-839 impact on flux, and reciprocal glutamine synthesis.

**Methods:** Glutaminolytic TNBC (HCC1806) and poorly glutaminolytic ER+ (MCF-7) xenograft mice with or without CB-839, underwent dynamic [^11^C]glutamine PET. HPLC quantified fractional radioactivity of [^11^C]glutamine, soluble metabolites ([^11^C]glutamate, [^11^C]CO_2_), and macromolecule-incorporated metabolites from blood and tumor. A four-tissue compartment model characterized *GLS1* activity (*k_GLS_*) and flux, *glutamine synthetase* activity (*k_GS_*), and subcellular glutamate distribution by comparing single vs. dual glutamate pool models. Averaged tumor curves and HPLC-derived tumor metabolites were fit. Monte Carlo simulations assessed parameter estimation performance.

**Results:** The single glutamate pool model showed high correlations between *k_GLS_* and other parameters, yielding inflated *k_GLS_* estimates. The dual glutamate pool model reduced correlations, improved *k_GLS_* recovery, and yielded subcellular glutamate distributions consistent with *in vitro* measurements. In TNBC, *k_GLS_* was 3-fold higher than ER+ tumors (non-overlapping 95% CI) with glutamate concentrated in the mitochondrial compartment. CB-839 reduced *k_GLS_* in TNBC and depleted mitochondrial glutamate (non-overlapping 95% CI), though glutaminolytic flux showed no distinguishable change. ER+ tumors showed higher *k_GS_* compared to TNBC.

**Conclusion:** [^11^C]glutamine PET kinetic analysis reveals distinct glutamine metabolic phenotypes in breast cancer subtypes. Preserved glutaminolytic flux and cytosolic glutamate in TNBC provide mechanistic hypotheses for clinical failure of *GLS1* inhibitors, informing ongoing studies.

## Introduction

“Glutamine addiction” is a phenotype observed in many aggressive tumors, including triple negative breast cancer (TNBC). These tumors show high glutamine dependence [1–3], mediated by glutaminolysis. Glutamine is converted to glutamate via *glutaminase* (*GLS1*), the rate-limiting step of this process [4], which can be followed by entry as α-ketoglutarate into tricarboxylic acid (TCA) cycle for ATP production or biosynthesis [3]. Consequently, *glutaminase* inhibition has emerged as a promising therapeutic strategy that is hypothesized to be effective in combination with other anti-cancer drugs. However, the kidney-type *glutaminase* enzyme (*GLS1*) inhibitor CB-839 (telaglenastat) showed disappointing anti-tumor efficacy in clinical trials [5, 6]. A reason for the poor efficacy is attributed to the inability to identify glutaminolytic tumors, highlighting a critical need for better selection markers to identify patients with this phenotype that are more likely to respond to targeted *GLS1* therapy. Recent work from our group suggests that dual inhibition of *GLS1* via CB-839 and glutamate transport can be an effective therapeutic strategy [7]. Glutamine positron emission tomography (PET) radiotracers provide a non-invasive approach to identify these markers through *in-vivo* assessment of glutamine metabolism and kinetics. In prior studies from our group, we demonstrated the ability to assess intracellular glutamine pool size, as an inverse indicator of *GLS1* activity, with ^18^F-(2*S*,4*R*)4-Fluoroglutamine ([^18^F]F-glutamine)[8, 9]; a minimally metabolized glutamine analog. Using *L*-[5-^11^C]-glutamine ([^11^C]glutamine), which is chemically identical to native glutamine, enables direct assessment of *GLS1* activity as well as downstream metabolic pathways. [^11^C]glutamine has been evaluated in preclinical and early human studies [10, 11]. In this effort, we probe the kinetics of *in-vivo* glutamine metabolism via [^11^C]glutamine and the impact on glutaminase inhibition with CB-839 for in vivo mouse models of breast cancer.

In breast cancer mouse models in [11], we showed that the rapid metabolism of [^11^C]glutamine into [^11^C]glutamate results in overlapping total time activity curves (TAC), where the relative contribution of parent tracer versus downstream metabolites to the total carbon-11 signal cannot be distinguished. This overlap makes it challenging to infer glutaminase activity from [^11^C]glutamine PET imaging alone. Key insights from [11] are as follows:

1. Tumor-specific metabolite profiles: glutaminolytic (HCC1806) tumors displayed high conversion to [¹¹C]glutamate while non-glutaminolytic (MCF-7) tumors showed lower conversion to glutamate and higher [¹¹C]glutamine fractions.
2. Glutaminase inhibition increased [^11^C]glutamine and reduced [^11^C]glutamate fractions in HCC1806 tumors.
3. A large [^11^C]glutamate pool in HCC1806 tumors originated primarily from intracellular glutamine catabolism rather than circulating [^11^C]glutamate.

We hypothesize that incorporating tumor metabolite measurements in the modeling process would enable [^11^C]glutamine PET flux quantification and guide mechanistic insights into therapeutic response. Here we present a compartment model describing [^11^C]glutamine kinetics in breast tumors, with parameterization aided by evidence from literature. This model and subsequent analysis examines three key components of glutamine metabolism from *in-vivo* data (**Figure 1A**):

1. glutaminolytic flux differences in response to glutaminase inhibitors, between tumors with high *GLS1* (HCC1806) and low *GLS1* (MCF-7) activity,
2. the reciprocal relationship of *glutamine synthetase* (*GS*) versus *glutaminase* (*GLS1*) in breast tumors,
3. the contribution of *GLS1*-derived glutamate versus xCT-transported extracellular glutamate to the cellular glutamate pool and the extent to which this pool is compartmentalized in the mitochondria [11–13].

**Figure 1:**
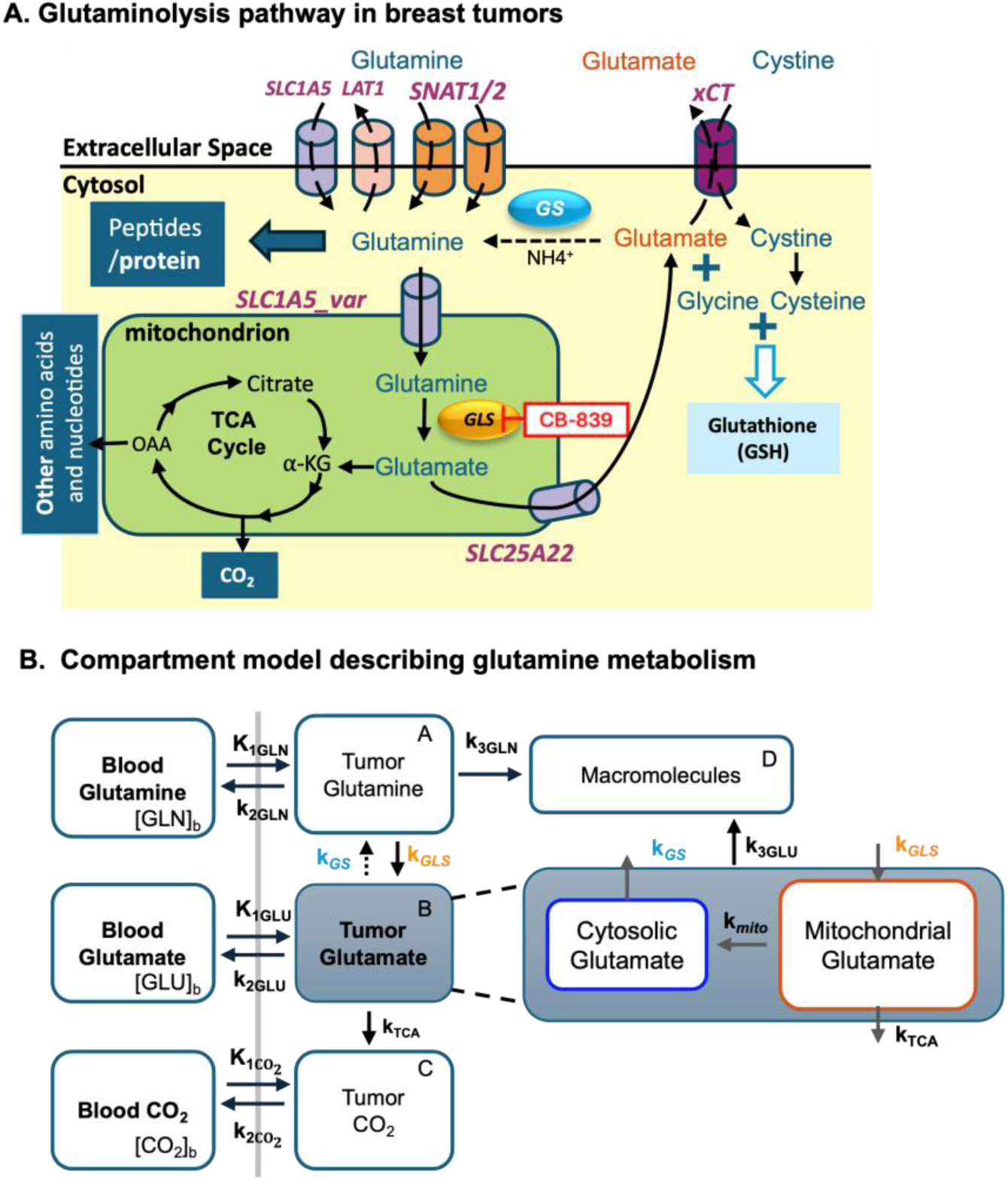
(A) Schematic showing pathway of glutamine metabolism in breast tumors as it enters the tumor and is metabolized as glutamate via *GLS1.* (B) Compartment model for [^11^C]glutamine. Subcellular compartmentation of glutamate is explored by modeling glutamate as a single pool versus separating into cytosolic and mitochondrial compartments

In this study, kinetic modeling of [^11^C]glutamine presents mechanistic insights to be tested experimentally in future studies and provides potential factors for why glutaminase inhibitors have not yet been effective in the clinic.

## Materials and Methods

### [^11^C]Glutamine PET studies and HPLC assay in Xenograft Tumors

Details of [^11^C]glutamine PET studies have been previously described [11] and are briefly summarized here. Mice with breast cancer xenograft tumors were dynamically imaged with [^11^C]glutamine over 30 minutes to acquire total radioactivity curves for blood and tumor. At specific intervals after tracer injection (t= 10, 20, 30 min p.i), ^11^C-labeled fractional activity of small soluble metabolites, macromolecules (e.g., peptides/proteins, referred to as pellet), and CO_2_ were measured in both blood and tumor. Fractional radioactivity of [^11^C]glutamine, [^11^C]glutamate and downstream metabolites ([^11^C]other) were measured via HPLC assay. Since the activity of soluble, non-CO_2_ downstream metabolites ([^11^C]other) comprised of less than 10% of total metabolite activity in the blood and tumor, it was omitted from further analyses. Details on the partial volume, delay, and metabolite corrections performed on the input functions and tumor TACs are included in [11]. The total tumor time activity curve (TAC) and tumor metabolite TACs were included in model fitting. See Figures 3 and 4 in *<u>Estimated metabolite-specific activity curves</u>* of our companion paper for more details regarding total and metabolite-specific TACs from blood and tumor [11].

### Compartment Model

The overarching framework of biochemical pathways for glutamine metabolism for evaluating glutamine kinetics is shown in **Figure 1A** and parameter definitions are included in **Table 1**. Rapid systemic metabolism of [^11^C]glutamine results in circulating labeled metabolites (i.e., [^11^C]glutamate, [^11^C]CO_2_) in plasma shortly after injection [11]. The tumor kinetics of [^11^C]glutamine, [^11^C]glutamate, and [^11^C]CO_2_ are described by three coupled compartment sets (A-C), each driven by their respective blood input functions (**Figure 1B**). The tumor [^11^C]glutamine compartment (Eq. 1, Figure 1B, A) represents the total intracellular glutamine pool, as glutamine uptake in breast cancer cells occurs through system ASC and system A transporters [14, 15]. *Glutaminase*-mediated conversion of glutamine to glutamate is the primary enzymatic step of glutaminolysis and principal parameter of interest in this model, as it differs substantially between tumor subtypes and a pharmacologic target of CB-839. The tumor [^11^C]glutamate compartment (Eq. 2, **Figure 1B**, B) represents glutamate derived from mitochondrial metabolism of glutamine via *GLS* and glutamate transported from interstitial space via *xCT* [13, 16]. The model also accounts for *de novo* glutamine synthesis from glutamate via *glutamine synthetase* [17]. The tumor [^11^C]CO_2_ compartment (Eq. 3, **Figure 1B**, C) represents CO_2_ generated from both glutamate-derived α-ketoglutarate oxidized in the TCA cycle and labeled CO_2_ /HCO_3_^-^ transported from blood [18, 19]. Lastly, the model also accounts for irreversible macromolecular trapping ([^11^C]macromolecules) (Eq. 4, Figure 1B, D) of glutamine and labeled downstream TCA metabolites that have been incorporated into biosynthetic products such as peptides, lipids and nucleic acids over the observation period [20, 21].

**Table 1:** [^11^C]glutamine model parameter ranges and starting values.

| Parameter | Description | Units | Minimum | Maximum | Initial Value |
| --- | --- | --- | --- | --- | --- |
| $K_{1\text{GLN}}$ | Glutamine blood-to-tissue transfer | mL/min/g | 0.01 | 0.2 | 0.067 |
| $K_1/k_{2\text{GLN}}$ | Glutamine partition coefficient | mL/g | 0.1 | 5 | 0.5 |
| $K_{1\text{GLU}}$ | Glutamate capillary transfer | mL/min/g | 0.001 | 0.1 | 0.005 |
| $K_1/k_{2\text{GLU}}$ | Glutamate partition coefficient | mL/g | 0.1 | 5 | 0.2 |
| $K_{1\text{CO}_2}$ | $\text{CO}_2$ capillary transfer | mL/min/g | $1 \times 10^{-7}$ | 0.2 | 0.002 |
| $K_1/k_{2\text{CO}_2}$ | $\text{CO}_2$ partition coefficient | mL/g | 0.2 | 1.2 | 0.5 |
| $k_{3,\text{GLN}}$ | Peptide incorporation rate constant | 1/min | 0.001 | 0.1 | 0.0119 |
| $k_{\text{GLS}}$ | Glutaminase rate constant | 1/min | 0.01 | 0.2 | 0.05 |
| $k_{\text{TCA}}$ | Rate constant of TCA turnover | 1/min | 0.001 | 0.2 | 0.05 |
| $k_{\text{GS}}$ | Glutamine synthetase rate constant | 1/min | 0.001 | 0.2 | 0.01 |
| $k_{3,\text{GLU}}$ | Peptide incorporation rate constant | 1/min | 0.00001 | 0.1 | 0.01 |
| $k_{\text{mito}}$ | Mitochondrial-to-cytosolic glutamate export rate constant | 1/min | 0.00001 | 0.1 | 0.005 |
| $V_b$ | Tissue blood volume | mL/g | - | - | 0.05* |

\*fixed value during optimization

Based on these kinetic compartments and using the notation in **Figure 1**, the following differential equations describe the compartment model. The tracer concentration is expressed in % injected activity per gram of tissue (%IA/g). Units and descriptions for all the variables are given in **Table 1**. We assume a tissue density of 1 g/mL.

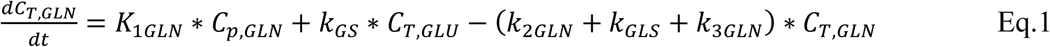

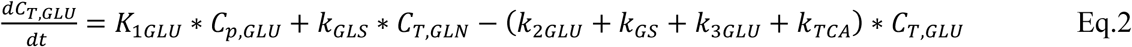

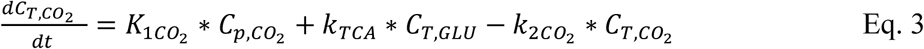

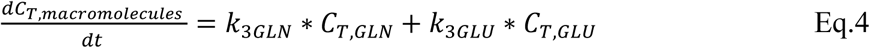

The total tumor activity, accounting for blood volume fraction that is recorded by the dynamic PET study is represented below:

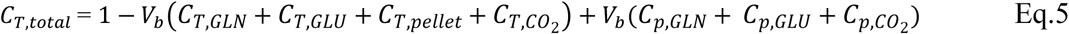

### Model Assumptions

#### Glutamine transport is reversible and limited by blood flow

Glutamine, the most abundant circulating amino acid (plasma: 0.5 – 0.8 mM [22]), is taken up by tumors at rates exceeding all other amino acids, with majority of intracellular glutamine typically originating from extracellular uptake in TNBC tumors [23]. Glutamine transport in solid tumors is mediated by group of amino acid transporters (system A and system ASC) [14, 24, 25] [26]. Michaelis-Menten transport kinetics showed Km values of 0.15 – 0.57 mM (in solid breast, colon and liver tumor derived cell lines), which is lower than plasma glutamine concentration and results in transporters operating at Vmax = 7.3 – 21 nmol/min/mg-protein [24]. Assuming a density 0.6 ml of cell water per tissue ml yields a transport capacity of 3-6 ml/min/g, exceeding tumor blood flow (0.09-0.2 ml/min/g) [27, 28]. Thus, glutamine blood-to-tissue transport rate constant (*K*_1*GLN*_) is assigned physiologic range of tumor blood flow: 0.01 – 0.2 ml/min/g (**Table 1**).

Intracellular glutamine is exported by LAT1, an obligatory antiporter [29], in exchange for essential amino acids like leucine. Since endogenous glutamine levels in breast cancer cells (0.3 mM) are well below the Km of LAT1 (2.2 mM) [30], first order kinetics are assumed. The glutamine efflux rate constant *k*_2*GLN*_, was derived as Vmax/Km (Vmax = 2.1 – 41.6 nmol/mg-protein/min [30, 31] assuming same cell water fraction), yielding *k*_2*GLN*_ bounds of 0.16 - 1.9 min^-1^. The substrate-limited efflux by LAT1 creates kinetic asymmetry with ASCT2, which operates at Vmax, resulting in measured glutamine influx-to-efflux ratios of 3-4-fold, favoring glutamine influx [29, 32]. The efflux rate constant was reparametrized as a ratio of glutamine uptake, as *K*_1*GLN*_ /*k*_2*GLN*_, ranging from 0.1–5 mL/g (**Table 1**).

We assume that, overall, intracellular glutamine behaves like a well-mixed precursor pool in cytosol and mitochondria, a simplification validated by rapid equilibrium kinetics observed in [^13^C]glutamine tracing studies [23]. The model assumes steady state concentration of native (i.e., unlabeled) molecules thus the content of tissue glutamine compartment (*V_TGLN_*) is considered constant and therefore can be described as:

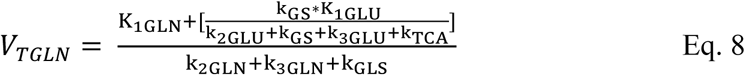

#### Tumor glutamate uptake is minimal, and intracellular glutamate is exchanged for cystine

TNBC and ER+ breast tumors overexpress the cystine-glutamate antiporter system Xc^-^(referred as xCT henceforth), which exchanges intracellular glutamate for cystine to support glutathione synthesis [33]. The Km for intracellular l-glutamate efflux (0.2 mM [34]) is substantially exceeded by the large intracellular glutamate pool (20 mM [34]), indicating that xCT operates near Vmax for glutamate export. The extracellular cystine concentration (0.028 mM [34]) is below Km for cystine influx (0.043 mM [35]), indicating that cystine import is substrate-limited. Together these conditions support net glutamate efflux via the xCT.

Although high affinity sodium-dependent transporters (system X^-^AG) cooperate with xCT to mediate glutamate influx in C6 glioma and lung cancer cells [36], it accounts for only 35% of total glutamate uptake in C6 glioma cells, and is comparatively less expressed in breast cancer [37, 38]. Glutamate transport in cultured fibroblasts is saturable with Km of 0.068 mM (extracellular glutamate concentration: 0.06 mM [39]) and thus we model transport by first-order kinetics, with Vmax of 0.68 nmol/mg/min with 100 mg-protein/g tissue, yielding glutamate blood-to-tissue transport rate constant (*K*_1*GLU*_) of 0.1 mL/min/g. Given the lower expression of glutamate transporters in tumors compared to healthy tissue, *K*_1*GLU*_ ranges between 0.001 – 0.1 mL/min/g (**Table 1**) Consistent with observations in lung cancer and glioma cell lines, where glutamine and leucine transport into the cell exceed that of glutamate [38], *K*_1_*_GLU_* is constrained to be less than *^K^*1*GLN*.

Using [^14^C]cystine tracing in xCT-expressing oocytes, Sato et. al confirmed 1:1 stoichiometry of cystine-glutamate exchange, and a glutamate efflux rate (48.5 – 75 pmol/10 min/oocyte) equal to cystine influx [40]. Based on oocyte cell volume (460 nl/oocyte) and in-vitro glutamate concentration (5.6 mM), this yields glutamate efflux rate constant bounds of 0.001 – 0.005 min^-1^. To reduce model complexity, glutamate efflux is reparametrized as ratio of glutamate uptake (K_1GLU_/k_2GLU_: 0.1 – 5 mL/g) (**Table 1**).

Despite glutamate transport via xCT, the large intracellular glutamate pool in glutaminolytic tumors is governed by glutamine uptake and elevated glutaminolysis [33]. We showed in preclinical studies that 60% of [^11^C]glutamine activity was retained as [^11^C]glutamate by 20 minutes, with minimal downstream metabolism to TCA intermediates [11]. This is supported by [^13^C]glutamine tracing in U87-MG cells, where cytosolic glutamate labeling reached 30% within 30 minutes and 55% within 120 minutes [38]. These studies support the observation that cancer cells do not significantly depend on extracellular glutamate to maintain their large and stable intracellular glutamate pool. At steady state, the distribution volume of tissue glutamate compartment (*V_TGLU_*) is described by:

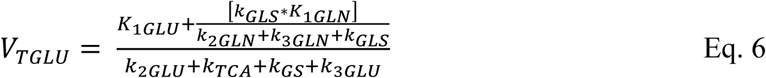

#### Glutaminase-directed glutamine metabolism is the rate-limiting step in glutaminolysis

*Glutaminase C* (GAC), a splice variant of *GLS1*, is a key *glutaminase* isozyme in several cancer cell lines [41], and is localized within the mitochondria. GAC shows superior phosphate-activated catalytic efficiency compared to other glutaminase isozymes, explaining its preferential tumor upregulation [42]. Mitochondrial glutamine transport via SLC1A5_var, is five-fold faster than the glutaminase reaction, and is thus excluded from modeling [43].

It has been shown that highly proliferative tumors such as TNBC and luminal B have higher glutamine metabolism and increased *GLS1* activity compared to poorly proliferative tumors [4], hence explaining the wide range of *GLS1* levels observed in breast cancer cell lines from 746.7 - 1380 nmol/mg/min (yielding estimates of 0.035-0.064 min^-1^, using in vitro glutamine concentration of 21375 nmol/mg-protein) [24]. Another study performed in cultured fibroblasts, estimated intracellular glutamine turnover rate to be 0.09 min^-1^ and observed that glutamine conversion to glutamate accounted for 25% of cell glutamine turnover rate [44]. Based on these observations, the expected range for k_GLS_ was set to 0.01 – 0.2 min^-1^ (**Table 1**).

Since *GLS1*-directed glutamine conversion to glutamate is considerably slower than subsequent enzymatic steps in glutaminolysis, *GLS1* serves as the primary regulator of glutaminolytic flux [24, 45]. This is further supported by several metabolic studies where *GLS1* inhibition dramatically reduced downstream TCA labeling and CO_2_ production [46, 47]. Reported glutaminolytic flux values range from 0.38 – 0.54 nmol/mg/min in murine B-lymphocyte hybridoma cells, Chinese hamster fibroblasts and kidney cells [45, 48, 49], and from 0.22 – 0.69 nmol/mg/min in breast cancer cells. Based on a tumor glutamine concentration of 0.3 mM and assuming 100-mg protein per tissue gram, Flux*_GLS_* bounds were set to 0.01– 0.23 mL/g/min, providing physiologically grounded reference range for modeling [26]. The first order dependence on the tumor glutamine compartment of tracer glutaminolysis, leads to a description of the rate constant of glutaminase activity (k_GLS_).

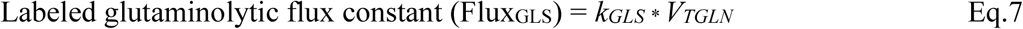

#### Glutamine and glutamate protein synthesis

The irreversible incorporation into macromolecules (with rate constant *k*_3*GLN*_) is a major possible fate of glutamine in proliferative tumors. Equilibration of intracellular glutamine/glutamate precursor pool occurs within minutes, suggesting that protein synthesis is a rate-limiting step with first order dependence on precursor pool specific activity [50]. This is directly evidenced by [5-^11^C]glutamine studies showing 30% protein incorporation at 30 minutes in glioma cells [10]. The first order equation is applied, resulting in *k*_3*GLN*_ equal to 0.0119 min^-1^ (plausible range between 0.001 – 0.1min^-1^ reflecting variability across tumor types) (**Table 1**) and further supported by 10% of total radioactivity trapped in macromolecular compartment in [^18^F](*2S,4R*)4F-glutamine studies [51].

The irreversible commitment of glutamine-derived glutamate to biosynthesis via *glutamate dehydrogenase* (GDH), is represented by *k*_3*GLU*_. Glutamate is converted into α-ketoglutarate, a regulated TCA cycle entry point. Although direct glutamate incorporation is minimal, as shown in [^18^F]-(2S,4R)-4F-glutamate studies [20], [3-^13^C]glutamine tracing showed that glutamine-derived glutamate provided 90% of anaplerotic oxaloacetate for lipid synthesis (nearly 25% of fatty acyl carbons) and rapid aspartate labeling for nucleotide biosynthesis [23]. Based on GDH catalytic efficiency (Vmax/Km = 0.008 min^-1^) in hepatomas [21], *k*_3*GLU*_ bounds were set to 0.00001 – 0.1 min^-1^ (**Table 1**), necessarily smaller than *k*_3*GLN*_.

#### Glutamine synthetase is primary source of de novo glutamine

*Glutamine synthetase* (*GS*) is assumed to be the main source of *de novo* glutamine when extracellular glutamine is limiting. This is supported by studies in hepatoma and neuroblastoma cells that show increased *GS* expression in response to glutamine deprivation [52, 53]. Upregulated *GS* activity is observed in well-differentiated tumors under metabolic stress to preclude futile cycling and maintain nitrogen balance [54, 55], and further supported by findings in primary mouse liver cells, where *GS* activity was four times greater than *glutaminase* activity [45]. Therefore, glutamine homeostasis in tumors is set by the balance between *glutaminase* and *GS*, and *GS* is a major control point in maintaining intracellular glutamine levels. *GS* follows Michaelis-Menten kinetics (glutamate Km = 3 mM [24]). In breast cancer cell lines, rate of *GS* ranged from 0.22 – 1 nmol/mg/min [24]. Assuming the same intracellular glutamate concentration and cell water fraction of tissue volume as earlier, maximum *GS* rate constant (*k_GS_*) is nearly 0.2 min^-1^, and *k_GS_* bounds are set to 0.001 – 0.2 min^-1^ (**Table 1**).

#### Systemic metabolism of [^11^C]glutamine is predominant [^11^C]CO_2_ source

Carbon dioxide (CO_2_) is assumed to be in a well-mixed pool that equilibrates rapidly according to tissue pH [56, 57]. The ^11^C-label at C5 of glutamine is retained through the first TCA turn via oxidative decarboxylation at α-ketoglutarate dehydrogenase, where unlabeled C1 is released as CO_2_ and ^11^C-label is incorporated into succinyl-coA. The label is released as [^11^C]CO_2_ in the second TCA turn when label reaches isocitrate dehydrogenase [23], meaning the CO_2_ compartment reflects cumulative TCA cycling. The net rate of ^11^C label transfer from glutamine to CO_2_ pool through this multi-step process is represented by TCA turnover rate constant (*k_TCA_*). With tumor glutamine pool ranging from 1– 5 nmol/mg-protein [51], and glutamine oxidation rate of 0.22 – 0.69 nmol/mg/min (for single carbon recovery) [3, 44, 45], *k_TCA_* is estimated to vary between 0.015 – 0.046 min^-1^. Given the significant variability in glutamine pools among tumor subtypes, the plausible range is broadened to 0.001 – 0.2 min^-1^ (**Table 1**).

Tumor-derived CO_2_ is highly membrane-permeable and rapidly mixes with a larger blood CO_2_ pool, derived mainly from liver and kidney (greater than 80% total CO^2^). This is consistent with findings from [^11^C]thymidine/[^11^C]CO_2_ modeling studies [18, 19]. Accordingly, [^11^C]CO_2_ is modeled here as a metabolite whose distribution volume is determined by pH-dependent bicarbonate buffering rather than metabolic trapping, with estimated distribution volume of 0.2 – 1.2 mL/g (**Table 1**) based on [^11^C]thymidine studies [18].

#### Subcellular glutamate compartmentation in breast tumors

Recent studies have proposed mitochondrial-to-cytosolic glutamate gradients resulting in efflux of intracellular glutamate through xCT for glutathione synthesis, while maintaining intracellular glutamate pool [12, 58, 59]. The mitochondrial pool is produced by *GLS1* activity, while the cytosolic pool is primarily fed by unidirectional export via SLC25A22 [58]. Net glutamate flux is directional from mitochondria to cytosol rather than fully equilibrated between compartments, allowing finite transport kinetics to be considered. However, the degree to which glutamate is exchanged between mitochondrial and cytosolic pools is an active area of investigation. Supporting compartmentalization of glutamate in distinct subcellular pools, [^15^N]glutamate incubation studies in fibroblasts showed that only a low level of extracellular label reached intracellular glutamate pool [44]. Thus, we examined in a secondary analysis whether explicitly separating cytosolic and mitochondrial glutamate pools meaningfully affects model behavior and kinetic parameters (**Figure 1A and B**). Mitochondria and cytosolic pools are treated as separate while total intracellular glutamate is assumed to be at quasi-steady over the short duration of tracer experiment. Direct subcellular fractionation measurements in glutaminolytic Hela cells showed that the total mitochondrial glutamate pool is approximately 25% of the cytosolic pool (log^10^(MT/CY) = −0.6). Accounting for the 10-fold smaller mitochondrial volume [60], the mitochondrial glutamate concentration is estimated to be 3-fold higher than cytosolic pool [16]. Based on *in vitro* observations of higher mitochondrial-to-cytosolic concentration gradients, a gaussian prior on the rate of mitochondrial glutamate export (*k_mito_*) was applied in model fitting and it was constrained between 0.00001-0.1 min^-1^ (**Table 1**). This prior-informed model is presented as hypothesis-generating to allow the PET data to dominate while guiding the rate towards physiologically plausible values. Thus, at equilibrium, the total glutamate volume of distribution (Eq. 9) can be considered as a combination of cytosolic (Eq. 10) and mitochondrial (Eq. 11) glutamate. Total tissue glutamine volume of distribution is described in Eq. 12.

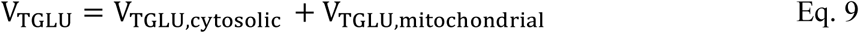

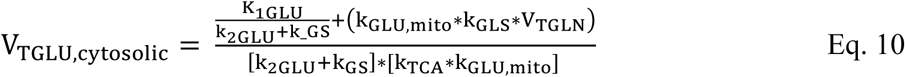

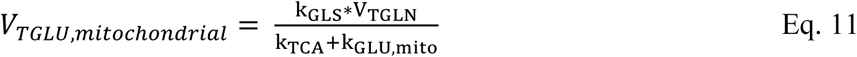

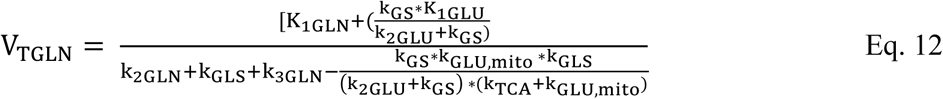

### Model Parameter Estimation and Simulations

The tumor and blood TACs were input into MATLAB. Image-derived arterial input functions served as model input. Fractional metabolite radioactivity in blood and tissue measured by HPLC at 10, 20, and 30 minutes post-injection, was interpolated over the PET acquisition interval to generate the soluble, pellet (macromolecule), glutamine, and glutamate TACs, as components of total tumor activity [11]. These data were used for model fitting to estimate the kinetic parameters described above.

#### Model Calibration and Kinetic Parameter Estimation

The compartment model was fit to averaged murine TACs per each of the two tumor and treatment types, using a global evolutionary optimization algorithm, (differential evolution toolbox, MATLAB [61, 62]). The optimization was configured with a population size of 100 and maximum of 100 iterations. The objective function minimized a weighted composite Huber loss, given its robustness to outliers, across total tumor activity and its components: soluble tissue, pellet, and metabolite-specific glutamine and glutamate TACs. Various weighting functions were evaluated (**Supplementary Figure 1**), and frame-duration weighting was applied in each component to emphasize later timepoints that are reflective of steady-state tracer distribution. Specifically, frame duration weights were computed as the ratio of each frame’s duration to mean frame duration, reflecting inverse relationship between frame duration and Poisson noise. These weights were normalized to preserve scale of loss function (weights summing to the number of frames). A discrete scaling was then applied to the final 3-4 timepoints for each component. A physiologic constraint penalty enforced that *K*_1*GLU*_ remained lower than *K*_1*GLN*_ via activation of a quadratic penalty term (Eq. 13) when the following constraint is violated:

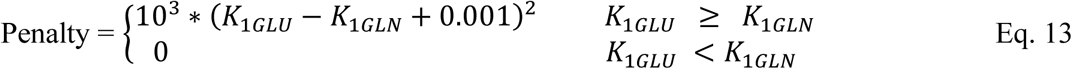

To prevent overfitting and stabilize parameter estimation, both lasso (L1 = 0.01) and ridge (L2 = 0.001) regularization terms were incorporated. The ODE system was solved using MATLAB’s ode45 solver.

#### Model Characterization: Sensitivity ahnd Identifiability Analysis

The total [^11^C]glutamine TAC cannot distinguish individual metabolite contributions [11], precluding reliable estimation of downstream kinetic parameters from total PET signal alone. To identify which metabolite-specific data provide additional experimental constraints necessary for reliable parameter estimation, variance-based global sensitivity analysis was performed. The Sobol’ method, uses analysis of variance decomposition [63, 64], to separate total output variance into first-order effects (SI: variance of each parameter alone) and total effects (ST: variance of each parameter including higher-order interactions) by sampling across the entire parameter space. Sensitivity indices were estimated from 1000 quasi-random parameter samples using Saltelli sampling scheme. Parameters with total effect index (ST) greater than 0.1 were considered sensitive [65]. Sensitivity was assessed independently for each measured data to characterize parameter sensitivity and overfitting risk.

Parameter identifiability was assessed through Fisher Information Matrix (FIM), by integrating the composite sensitivity matrices across all measured data. The Moore-Penrose pseudoinverse was used to compute parameter covariance and correlation matrices from singular value decomposition of sensitivity matrix [66].

Profile likelihood analysis was additionally performed for selected parameter pairs (*k_mito_* and *k_TCA_*) confirming practical identifiability and characterize collinear pairs. In the dual pool model, given the lack of sensitivity of the mitochondrial-to-cytosolic glutamate export parameter, *k_mito_* to any measured data (**Figure 4**), all remaining parameters were refit at each fixed *k_mito_* value. This revealed collinearity between the TCA turnover rate constant (*k_TCA_*) and *k_mito_* which jointly govern clearance from mitochondrial glutamate pool (**Supplementary Figure 3B,D**). *k_TCA_* was fixed at discrete values and all remaining parameters were refit. Cytosolic and mitochondrial glutamate concentration ratios at 30-minutes were highly sensitive to *k_TCA_*, with progressively unrealistic ratios (e.g. ratio ∼22 in MCF-7 vehicle). Since *k_TCA_*and *k_mito_*were not jointly identifiable, *k_TCA_* was fixed to 0.03 min^-1^ based on physiologic estimate derived from ^13^C MRS measurements (**Supplementary Figure 3C,E**).

#### Assessment of Parameter Estimation Performance using Simulated Curves with Added Noise

To assess kinetic parameter local identifiability, we performed Monte-Carlo simulations. Ground truth TACs were generated by calibrating model to group-averaged experimental data. Radioactive decay and frame duration were applied to the TACs to calculate tumor counts-representative of Poisson distribution of radioactivity. Gaussian noise was added to generate 250 noise realizations (μ = 0, σ = 1). The global scaling factor was empirically determined as the value that minimized the weighted sum of squared difference between the standard deviation of the synthetic TACs and experimental inter-subject variability across all timepoints [67, 68]. Under these conditions, the 25 to 30-minute data point (last frame) in the simulated TAC had a coefficient of variation of approximately 6-8%, depending on tumor type. The noise model was applied consistently to the total tumor and its derived soluble, pellet, glutamine and glutamate-specific TACs. The noisy curves were fitted in MATLAB using the corresponding compartment model, and the resulting parameter estimates are reported as median [2.5^th^, 97.5^th^ percentile] with 95% confidence intervals. For each true kinetic parameter 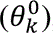 (*θ*^0^), the bias and precision (%SD) of kinetic parameter estimate 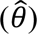 was also calculated (Eq. 14 and Eq. 15):

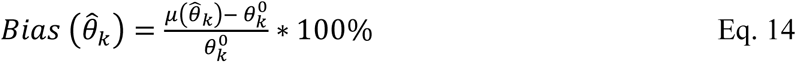

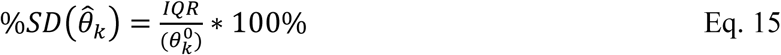

#### Statistical Analysis

Pairwise group comparisons were performed using non-overlapping 95% confidence intervals as the primary criterion for statistical significance. Model selection between single-pool and two-pool glutamate compartment models was assessed using AIC and BIC computed from unweighted residuals.

## Results

### Model Sensitivity Analysis

Global sensitivity analysis using Sobol indices was performed to quantify the information content of each measured data with respect to model parameters. Total effect indices (ST) for each parameter across all measured data are shown in **Figure 2**.

**Figure 2:**
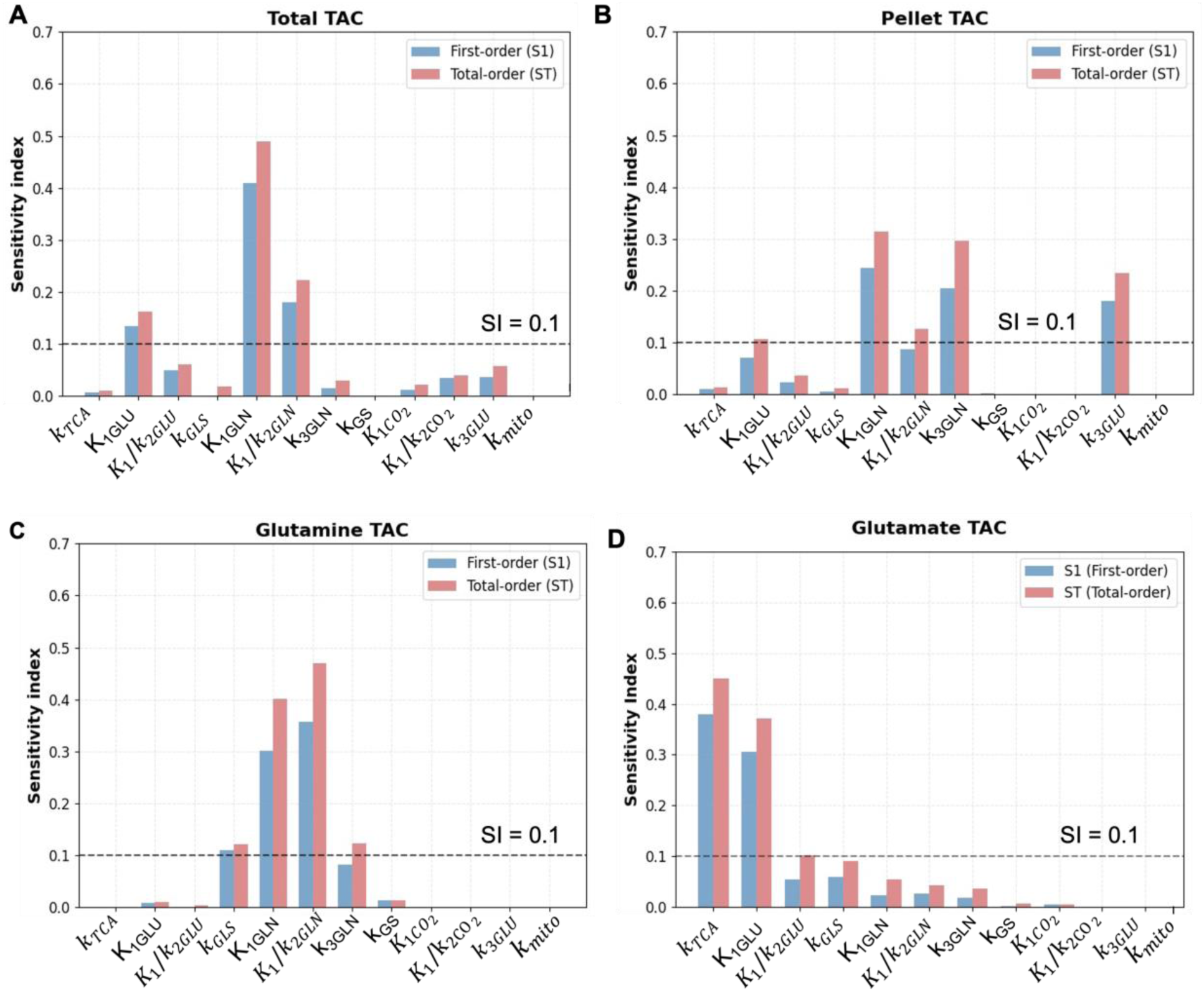
Sobol sensitivity analysis for (**A**) total tumor TAC, (**B**) pellet TAC, and individual metabolite TAC, including (**C**) glutamine TAC and (**D**) glutamate TAC

The total tumor TAC showed highest sensitivity to glutamine and glutamate transport parameters (*K*_1*GLN*_ ST=0.41, *K*_1*GLN*_ /*k*_2*GLN*_ ST = 0.18 and *K*_1*GLU*_ ST=0.13) (**Figure 2A**). The glutamine-specific TAC showed sensitivity to *k_GLS_* (ST=0.11) (**Figure 2C**). The glutamate-specific TAC showed highest sensitivity for *K*_1*GLU*_/*k*_2*GLU*_ (ST=0.1) and *k_TCA_* (ST=0.45) (**Figure 2D**). The pellet fraction showed highest sensitivity for *k*_3_*_GLN_* (ST=0.31) and low sensitivity to other parameters (**Figure 2B**). Parameters such as *k_GS_*, *K*_1*CO*2_, *K*_1*CO*2/_*k*_2*CO*2_ showed total effect indices below 0.1 across all measured data. Overall, *k*_3*GLN*_, *k_GLS_* and *k_TCA_* showed near zero ST in total tumor TAC but non-negligible indices in at least one metabolite-specific data.

### Compartment Model Calibration to Xenograft Data

The averaged TACs (soluble tissue extract, [^11^C]glutamine, [^11^C]glutamate) for TNBC vehicle (**Figure 3A and 3B**) and ER+ vehicle (**Figure 3C and 3D**) tumors were fit to the compartment model, with and without *glutamine synthetase* rate constant (*k_GS_*). The blood fraction was fixed at 5% based on [^18^F]4F-Gln breast cancer xenograft studies [8].

**Figure 3:**
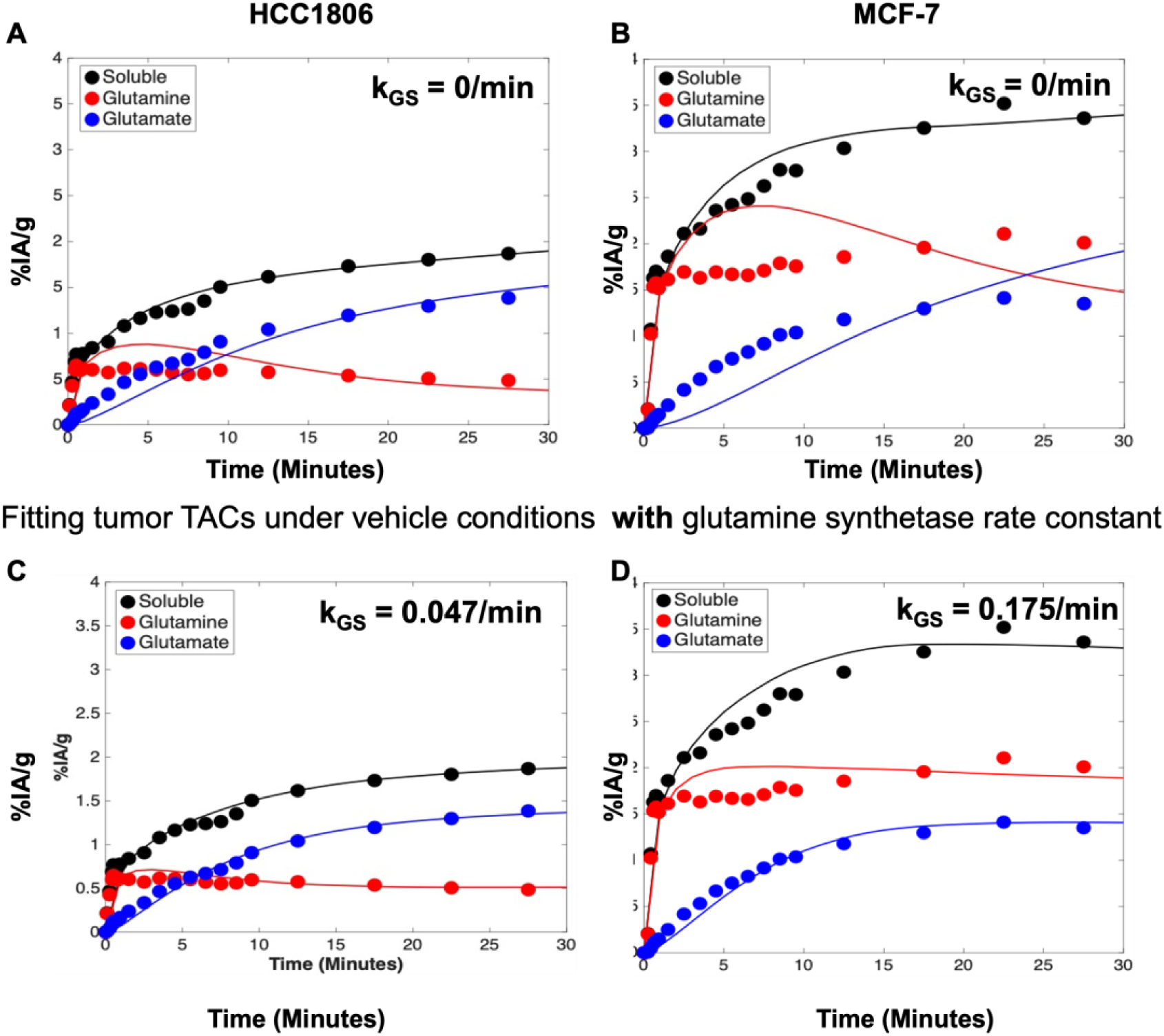
(A) and (C) show fitting of total tumor TACs and individual metabolite compartments for HCC1806 tumors with (A) and without (C) glutamine synthetase rate constant in the compartment model. (B) and (D) show MCF-7 tumor TACs and metabolite compartment model fitting with (B) and without (D) glutamine synthetase rate constant

The compartment model predicted the soluble tissue extract curve well in all the comparisons. However, the model fits to [^11^C]glutamine and [^11^C]glutamate activity varied based on tumor type. When *k_GS_* was 0 in TNBC vehicle tumors, the compartment model fitted qualitatively well with some overestimation at early timepoints, and model fit was improved by estimating *k_GS_* to be 0.047/min. In contrast, for ER+ tumors, the model without *k_GS_* fit poorly and incorrectly showed sharp decline in [^11^C]glutamine and equivalent rise in [^11^C]glutamate. By estimating *k_GS_* in the model at 0.175/min, the model fit to data was improved, suggesting that glutamine synthetase is a significant contributor of the glutamine pool in well-differentiated ER+ tumors, as has been noted in prior biologic studies [69].

### Model Characterization and Identifiability Analysis

The single glutamate pool model and dual glutamate pool models showed statistically indistinguishable goodness-of-fit across all experimental groups (ΔAIC=0.66-1.74; **Supplementary Table 2**). In the single glutamate pool model, parameter correlation matrix identified high correlations between *k_GLS_* (|r| ≥ 0.6) and *K*_1*GLN*_, K_1CO2_ and k_TCA._ Also *k*_3*GLN*_ was highly correlated with K_1_/k_2GLN_, K_1GLU_, K_1CO2_. The single pool formulation cannot distinguish glutamine metabolism from TCA cycle kinetics, and in turn the model finds inflated *k_GLS_* solutions (**Figure 4A-B**). In the dual glutamate pool model, these correlations were substantially reduced indicating improved structural identifiability for *k_GLS_* and k_3GLN_. A high negative correlation (r=-0.95) was retained between k_GLS_ and k_3GLN_ in dual glutamate pool model (**Figure 4**).

**Figure 4:**
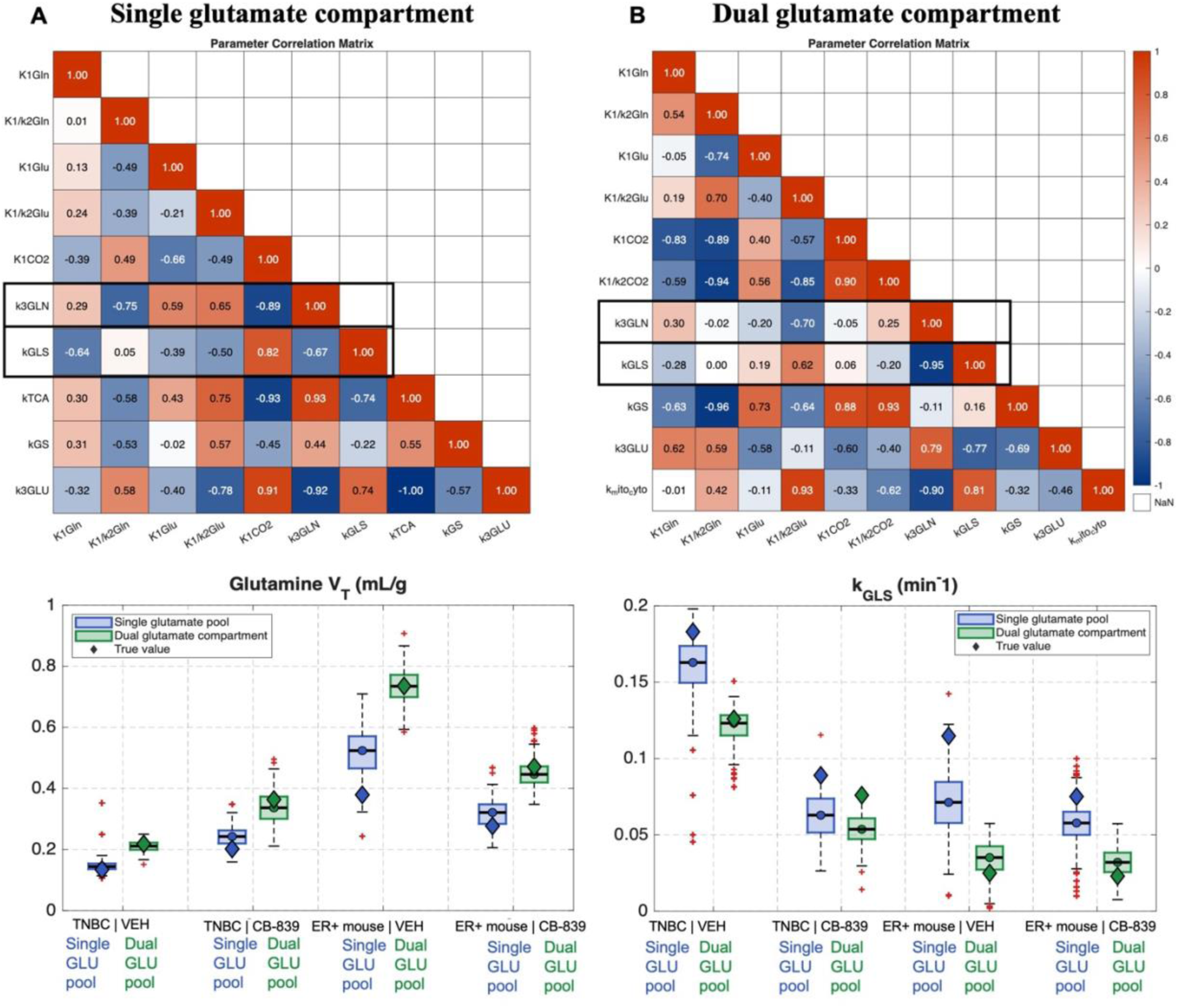
Parameter correlation matrices for (**A**) single glutamate pool model versus (B) dual glutamate pool model are shown for all the free parameters (k_TCA_ is fixed in dual glutamate pool model) for vehicle-treated HCC1806 tumors. The bottom row shows the comparison of estimation of V_TGLN_ (**C**) and k_GLS_ (**D**) from single glutamate pool and dual glutamate pool models after adding poisson noise to time activity curves and fitting each noise realization with corresponding model. Boxes reflect 25^th^, 50^th^, and 75^th^ percentiles of data, and whiskers are extent of data after outlier removal (1.5 × interquartile range, where interquartile range = 75th–25th percentiles). Diamonds show true V_T_ before noise is added to fit curve.

Monte Carlo simulations showed improved accuracy and precision of recovering the true glutamine *V_T_* and *k_GLS_* with the dual glutamate pool model compared to single glutamate pool, especially for MCF-7 tumors (**Figure 4C-D**). For glutamine *V_T_* the bias and precision were both improved in the dual glutamate pool model (bias, 0.1-8% vs. 8-38%; %SD, 10-20% vs. 13-28%, respectively). For *k_GLS_*, 95% confidence intervals were consistently narrower in the dual glutamate pool model (reduction of 16-45% across groups) despite comparable precision (%SD, 28.5% vs. 29.8%, respectively). Together with the statistically indistinguishable goodness of fit, improved parameter correlation structure, and physiologic consistency with subcellular compartmentalization of glutamate metabolism in cancer cells, these results support selection of dual glutamate pool compartment model for subsequent kinetic characterization. Single pool model parameter estimates are additionally provided in **Supplementary Table 3**.

### Subcellular Glutamate Distribution in Mouse Xenografts

The dual glutamate pool model explicitly partitions glutamate into cytosolic and mitochondrial compartments, enabling quantification of subcellular glutamate distribution volumes alongside kinetic parameters. Simulations showed that the glutamine blood-to-tissue transfer (K_1GLN_), glutamine volume of distribution (V_TGLN_), glutamate distribution volume (V_TGLU_), and macromolecular trapping rate constant (k_3GLN_) could be accurately and reliably recovered from the dual glutamate pool model (**Supplementary Table 4**). Glutaminase activity (*k_GLS_*) was recovered with acceptable accuracy and precision (mean bias, 12%; %SD, 35%), while *Flux_GLS_* showed comparable bias but wide confidence intervals (mean bias, 8%; %SD, 37%).

In TNBC vehicle tumors, dual glutamate compartment modeling recovered mitochondria-dominant glutamate distribution with 76% of total glutamate in the mitochondrial compartment (*V_TGLU_*_,*mitoc*ℎ*ondrial*_ = 0.51 mL/g, 95% CI [0.36-0.58]) and 22% in the cytosol (*V_TGLU_*_,*cytosolic*_: 0.15 mL/g, 95% CI [0.09 - 0.3]) (**Figure 5A-C**). This distribution is consistent with independent biochemical measurements: subcellular fractionation studies in Hela cells, reported a 3-fold higher mitochondrial concentration of TCA cycle intermediates [16, 70] and digitonin permeabilization in TNBC cells reported a 4-5-fold mitochondrial-to-cytosolic glutamate concentration gradient in vitro [59, 71].

**Figure 5:**
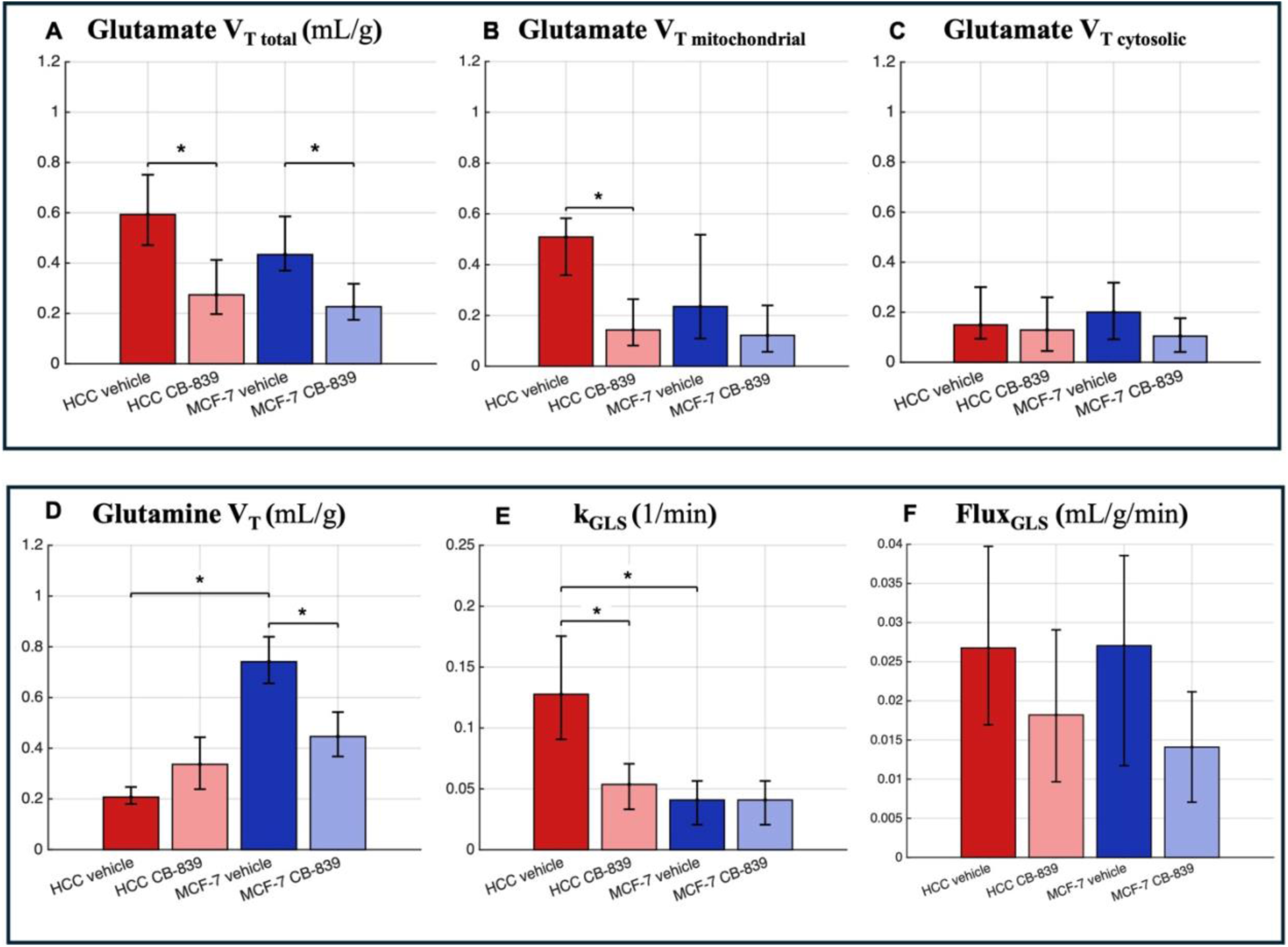
Kinetic parameter estimates are reported from noise realizations generated from poisson noise added to averaged time activity curves for each group of mice and fitting each realization with dual glutamate pool compartment model (**A-C**) median and interquartile range [2.5^th^ and 97.5^th^ percentile] is shown for total and compartmentalized (mitochondrial and cytosolic, respectively) glutamate volume of distribution. (**D-F**) Glutaminase flux is calculated based on glutaminase activity*glutamine volume of distribution. Error bars represent 95% confidence intervals (CI) from Monte Carlo simulations (N=250). Statistical significance (*) is determined by non-overlapping CI between groups.

The *glutaminase* activity in TNBC (*k_GLS_*: 0.12 min^-1^, 95% CI [0.09 - 0.14]) was 3.2-fold higher than ER+ tumors (*k_GLS_*: 0.037 min^-1^, 95% CI [0.016-0.05]; non-overlapping CI, **Figure D**). The high *k_GLS_* is expected to load the mitochondrial glutamate pool faster than it can be exported to cytosol [58]. Despite this high metabolic activity, TNBC showed relatively modest glutamine uptake (*K*_1*GLN*_: 0.028 mL/min/g, 95% CI [0.024 – 0.032]) and distribution volume (*V_TGLN_*: 0.21 mL/g, 95% CI [0.18-0.24]) (**Supplementary Figure 4B**, **Figure 5D**), indicating efficient intracellular conversion of imported glutamine. Glutamine macromolecular incorporation was also higher in TNBC compared to ER+ vehicle groups (*k*_3*GLN*_: 0.041 min^-1^, 95% CI [0.038-0.044] vs. 0.029 min^-1^, 95% CI [0.021-0.032]; non-overlapping CI, **Supplementary Figure 4**), consistent with elevated biosynthetic demand.

In contrast, ER+ vehicle tumors showed a balanced subcellular glutamate distribution (51% mitochondrial, 0.22 mL/g 95% CI [0.11-0.45], 49% cytosolic, 0.21 mL/g 95% CI [0.11-0.32] **Figure 5B-C**), consistent with absence of mitochondrial concentration gradient observed in digitonin permeabilization studies of ER+ cells [59, 71]. ER+ tumors showed lower *k_GLS_* (0.037 min^-1^, 95% CI [0.016-0.05]), but markedly higher glutamine uptake (*K*_1*GLN*_: 0.065 mL/min/g, 95% CI [0.054–0.074], which is nearly 2.3-fold higher than TNBC vehicle (*K*_1*GLN*_: 0.028 mL/min/g, 95% CI [0.024 – 0.032]) and higher glutamine synthetase (*k_GS_*, 0.105 min^-1^ resulting in a 3.5-fold larger glutamine pool (*V_TGLN_*: 0.74 mL/min/g, 95% CI [0.66-0.84]) (**Figure 5D-F**) compared to TNBC. Notably, *Flux_GLS_* in ER+ tumors (0.027 mL/min/g [0.012-0.039]) was comparable to TNBC tumors (0.026 mL/min/g 95% CI [0.018-0.03]) despite lower *k_GLS_*. The wide confidence intervals across experimental groups suggest that *Flux_GLS_* is not identifiable from [^11^C]glutamine studies.

### Estimates of Tumor Response to Glutaminase Inhibition

CB-839 treatment produced subtype-specific pharmacodynamic responses with mitochondrial-specific glutamate depletion in TNBC and reduced glutamine uptake and pool size in ER+ tumors. Assuming Michaelis-Menten first order kinetics [4, 72], *k_GLS_* is directly comparable to the GLS enzymatic rate. TNBC tumors treated with CB-839 show reduced glutaminase activity compared to vehicle group (*k_GLS_*, CB-839: 0.05 min^-1^, 95% CI [0.03-0.07] vs. vehicle: 0.12 min^-1^, 95% CI [0.098-0.14]; non-overlapping CI) (**Figure 5D-F**) accompanied by reduction in total glutamate distribution volume (*V_TGLU_*, CB-839: 0.29 mL/g, 95% CI [0.2-0.4] vs. vehicle: 0.67 mL/g, 95% CI [0.52-0.78]; non-overlapping CI, **Figure 5A**). This reduction was driven by depletion in mitochondrial pool (*V_TGLU_*_,*mitoc*ℎ*ondrial*_, CB-839: 0.14 mL/g, 95% CI [0.08-0.26] vs. vehicle: 0.51 mL/g, 95% CI [0.36-0.58]; non-overlapping CI, **Figure 5B**) while cytosolic glutamate showed overlapping 95% CI between treatment groups. *V_TGLN_* increased with CB-839 (CB-839: 0.34 mL/g, 95% CI [0.24-0.44] vs. vehicle: 0.21 mL/g, 95% CI [0.18-0.24], consistent with [^18^F]F-glutamine studies [8, 51]. *Flux_GLS_* showed overlapping 95% CI between CB-839 and vehicle groups (0.026 mL/min/g 95% CI [0.018-0.03] vs. 0.018 mL/min/g 95% CI [0.01-0.029], **Figure 5F**), indicating no distinguishable treatment effect on flux with CB-839.

In ER+ tumors, *k_GLS_* was not different between CB-839 and vehicle group (CB-839: 0.032 min^-1^, 95% CI [0.015-0.05] vs. vehicle: 0.037 min^-1^, 95% CI [0.016-0.05], **Figure 5E**), which is consistent with the low baseline *GLS* activity. *V_TGLU_* was reduced with CB-839 (CB-839: 0.23 mL/g 95% CI [0.17-0.32] vs. 0.43 mL/g 95% CI [0.37-0.59]; non-overlapping CI) although unlike TNBC, this reduction was not clearly attributable to either subcellular compartment. However, *V_TGLN_* (0.74 mL/g, 95% CI [0.66-0.84] vs. 0.45 mL/g, 95% CI [0.37-0.54] and *K*_1*GLN*_ (0.065 mL/min/g, 95% CI [0.054-0.074] vs. 0.037 mL/min/g, 95% CI [0.03-0.043]) were both reduced with CB-839. CB-839-directed *V_TGLN_* reduction may reflect *GS*-mediated glutamine recycling.

## Discussion

This study presents a compartment model for kinetic analysis of [^11^C]glutamine in mouse breast cancer xenografts integrating tumor metabolite-specific radioactivity (including large molecules in pellet and small molecule metabolites in soluble extract) as a pre-clinical tool for generating mechanistic hypotheses about glutamine metabolism and tumor response to *GLS1* inhibition. Previous work showed that total tumor activity and tumor-to-blood ratios showed only modest changes with *GLS1* inhibition due to the formation and systemic circulation of [^11^C]glutamate [11], motivating the inclusion of metabolite-specific measurements in kinetic modeling. Global sensitivity analysis showed that the total TAC as an input was insensitive to *k_GLS_* and *k_TCA_*. These parameters could only be estimated with inclusion of measured blood and tumor metabolite TACs for glutamine and glutamate, respectively, while the pellet fraction informed estimation of *k*_3*GLN*_. All parameters were retained as free parameters in model fitting, due to compartment interdependencies.

The tissue metabolites were initially represented with a single compartment per species. The single glutamate pool model yielded high parameter correlations between *k_GLS_* and *K*_1*GLN*_,*K*_1*CO*2_ and *k_TCA_*, and produced *k_GLS_* estimates that were 2-fold higher than reported enzymatic activity [24, 73]. Explicit separation of the cytosolic and mitochondrial compartments with the dual glutamate pool model reduced these correlations, and simulations showed improved accuracy and precision in recovering *k_GLS_* and *V_TGLN_* relative to single glutamate pool model. The dual glutamate pool model also produced subcellular glutamate distribution ratios consistent with direct biochemical measurements [70, 71]. On this basis, dual glutamate pool model was selected for subsequent kinetic analysis.

We previously showed that metabolite-specific TACs were concordant with other studies of glutamine metabolism [8, 11] with higher [^11^C]glutamate accumulation in glutaminolytic TNBC than ER+ tumors, and [^11^C]glutamate crossing over [^11^C]glutamine at approximately 10-minutes. This crossover, observed only in vehicle treated TNBC, was reproduced by the model from independently fitted parameters, consistent with the fitted *k_GLS_*.

In TNBC, the dual glutamate pool model recovered a larger mitochondrial glutamate volume of distribution relative to cytosol, consistent with the mitochondria-dominant glutamate distribution *in vitro* [38, 71]. This distribution may relate to the 3-fold higher *k_GLS_* in TNBC compared to ER+ tumors, consistent with reported differences in *GLS* enzyme kinetics [13, 69].

The elevated *GLS* activity operating on rapidly turning over glutamine pool in TNBC supports the mitochondria-dominant glutamate phenotype. In contrast, ER+ tumors showed no difference in subcellular glutamate distribution volumes, consistent with a distinct metabolic profile characterized by low *k_GLS_* and high *k_GS_*. *V_TGLN_* was larger in ER+ tumors, which may reflect intracellular glutamine accumulation supported by high ASCT2-mediated transport and *k_GS_* - mediated glutamine recycling. Together these observations suggest a glutamine independent phenotype of ER+ tumors [69].

Glutaminase inhibition with CB-839 produced distinct subtype-specific responses. In TNBC, CB-839 significantly reduced *k_GLS_* and *V_TGLU_* (40% of vehicle; non-overlapping CI), consistent with reported reduction in glutamate pool size to 33% of vehicle by *ex vivo* ^1^H MRS measurements [74]. This reduction was compartment specific: mitochondrial glutamate was reduce while cytosolic distribution volume showed overlapping CI between groups, consistent with digitonin-permeabilized subcellular fractionation studies [71]. The preferential depletion of the mitochondrial pool with CB-839 in TNBC adds kinetic evidence supporting the mitochondrial glutamate “buffer” hypothesis, in which the mitochondrial pool may serve as a reservoir sustaining cytosolic glutamate availability for supporting cystine import, rather than a freely-exchanging equilibrium pool. Conversely, the relatively preserved cytosolic distribution volume with CB-839, offers one hypothesis for why CB-839 alone may be insufficient to reduce cellular anti-oxidant capacity. This is consistent with our work showing that dual inhibition of mitochondrial and cytosolic glutamate sources (via erastin) may be required to deplete glutathione and generate sufficient oxidative stress [7]. *V_TGLN_* increased nearly 1.6-fold with CB-839 in TNBC. Given the reported 2.5-fold increase in plasma glutamine [51], the imaging-estimated glutamine pool size increase is approximately 1.6 ∗ 2.5 = 4-fold, which is congruent with the 4.3-fold increase in glutamine pool size from MRS with CB-839 [51]. Despite substantial reduction in *k_GLS_*, *Flux_GLS_* was decreased minimally (depicting overlapping 95% CI). This limitation arises from the compensatory relationship between *k_GLS_* and *V_TGLN_*, and gives insight into potential factor contributing to limiting clinical efficacy of CB-839. Independent measurements of glutamine and glutamate kinetics via [^13^C]glutamine stable isotope tracing studies will enable distinction of *Flux_GLS_*. In ER+ tumors, CB-839 produced no distinguishable change in *k_GLS_* (overlapping 95% CI) but reduced *V_TGLN_*, which may reflect decrease in *k_GS_* -mediated glutamine recycling with depletion of *k_GS_* substrate. This recycling path is not captured by [^18^F]F-glutamine studies, which contrastingly reported stable glutamine pool size with CB-839 [8, 51].

Several limitations of this study are acknowledged: Firstly, parameter estimation was performed on group-averaged TACs rather than individual mouse curves, limiting direct characterization of inter-subject variability. Second, though this study used two representative breast cancer xenograft models, with high and low *GLS*, future studies should use additional cell lines to validate the modeling framework. Third, this study did not directly measure the different sources of tumor metabolites, for example degree to which [^11^C]glutamate was derived from plasma versus intratumoral conversion, hence the compartment-specific glutamate distributions are intended to be hypothesis generating results constrained by specific physiologic assumptions. This is an active area of investigation to confirm the measurements of mitochondrial-to-cytosolic ratios through [^13^C]glutamine stable isotope tracing. We note that these are “plausibility” studies of proposed compartment model to generate hypotheses.

## Conclusions

The present study provides a framework for leveraging preclinical [^11^C]glutamine studies for developing and testing mechanistic hypotheses about glutamine metabolism in breast tumors. Integration of metabolite-specific radioactivity measurements enables estimation of *k_GLS_* and subcellular glutamate compartmentation that is not accessible from total tumor PET signal, suggesting that [^11^C]glutamine of limited utility in human studies. The primary methodological contribution form this study that leverage the ability to measure metabolites in both blood and tumor in mouse models to inform kinetic analysis, is that the dual glutamate pool model (separate mitochondrial and cytosolic pools) helps explain fundamentally distinct glutamine metabolic phenotypes observed for TNBC versus ER+ breast cancer. TNBC tumors exhibit high *GLS* activity driving mitochondria-to-cytosolic glutamate concentration gradients, with CB-839 ablating mitochondrial glutamate while cytosolic pool is unchanged rendering it vulnerable to *GLS* inhibition. ER+ tumors exhibit low *GLS* activity, balanced subcellular glutamate distributions, active *GS*-mediated glutamine recycling, rendering it less susceptible to *GLS* inhibition. These hypothesis-generating insight suggest future mechanistic studies to clarify the cellular biology of glutamine metabolism in cancer cells.

## Acknowledgement

We thank Calithera Biosciences Inc. for providing CB-839 and Dr. Marina Gelman for insightful discussions. Conception and study design: D.M, E.J.L, R.Z, A.RP, C.T.H, R.A.D Data curation: C.T.H, H.C., P.P, Data analysis and interpretation: R.A.D, E.J.L, D.M, C.T.H Writing–drafting the article: R.A.D, C.T.H, D.M, E.J.L, Writing—critical review and editing: C.T.H, D.M, E.J.L, R.Z. All authors have read and agreed to the published version of the manuscript.

## Data Statement

Data described in the manuscript, code book, and analytic code will be made available upon request pending application and approval

## Declaration of Generative AI and AI-assisted technologies in the writing process

During the preparation of this manuscript, the author(s) used Claude 3.5 Sonnet (Anthropic PBC, San Francisco, CA, USA) to improve the clarity, syntax, and grammatical flow of the text. After using this tool, the author(s) reviewed and edited the content as needed and take(s) full responsibility for the accuracy and integrity of the final publication.

