## Supplementary Figures for "*L*-5-[11C]-glutamine PET of Breast Cancer: Kinetic Analysis in Mouse Models to Evaluate Glutamine Metabolism"

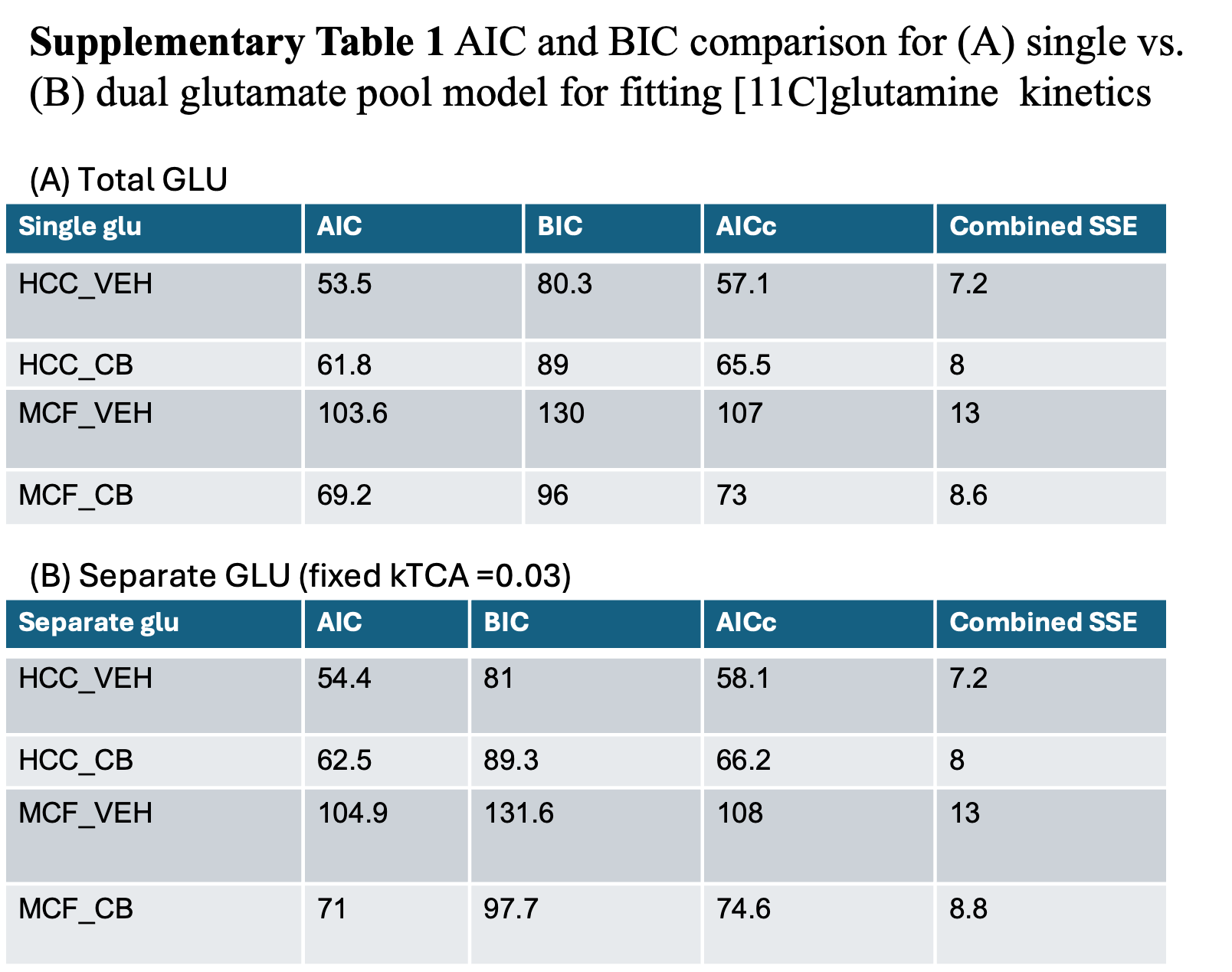

| **Supplementary Table 2. Kinetic parameter estimates for single glutamate pool model** | | | | |
| --- | --- | --- | --- | --- |
| *Values reported as median [2.5th – 97.5th percentile] from Monte Carlo simulations (n=500). Dashes (—) indicate parameter not included in single-pool model.* | | | | |
| **Parameter** | **TNBC Vehicle** | **TNBC CB-839** | **ER+ Vehicle** | **ER+ CB-839** |
| ***Rate Constants*** | | | | |
| K₁GLN (mL/min/g) | 0.028 [0.024 – 0.067] | 0.023 [0.019 – 0.067] | 0.058 [0.048 – 0.069] | 0.035 [0.030 – 0.067] |
| K₁/k₂GLN (mL/g) | 0.532 [0.269 – 0.982] | 0.659 [0.427 – 1.053] | 1.138 [0.586 – 1.690] | 0.720 [0.464 – 1.122] |
| K₁GLU (mL/min/g) | 0.018 [0.003 – 0.029] | 0.015 [0.000 – 0.023] | 0.035 [0.003 – 0.055] | 0.012 [0.001 – 0.032] |
| K₁/k₂GLU (mL/g) | 0.717 [0.200 – 1.578] | 0.383 [0.109 – 1.170] | 0.767 [0.266 – 1.506] | 0.367 [0.111 – 0.899] |
| K₁CO₂ (mL/min/g) | 0.079 [0.001 – 0.172] | 0.106 [0.002 – 0.181] | 0.049 [0.001 – 0.175] | 0.066 [0.002 – 0.159] |
| K₁/k₂CO₂ (mL/g) | 0.323 [0.243 – 0.519] | 0.695 [0.500 – 1.046] | 0.105 [0.023 – 0.489] | 0.383 [0.249 – 0.691] |
| k₃GLN (min⁻¹) | 0.041 [0.012 – 0.045] | 0.016 [0.012 – 0.021] | 0.023 [0.019 – 0.033] | 0.021 [0.012 – 0.022] |
| kGLS (min⁻¹) | 0.163 [0.050 – 0.195] | 0.063 [0.037 – 0.088] | 0.071 [0.033 – 0.119] | 0.058 [0.027 – 0.088] |
| kTCA (min⁻¹) | 0.017 [0.000 – 0.051] | 0.039 [0.010 – 0.091] | 0.011 [0.000 – 0.092] | 0.046 [0.001 – 0.125] |
| kGS (min⁻¹) | 0.051 [0.032 – 0.074] | 0.052 [0.000 – 0.095] | 0.105 [0.032 – 0.171] | 0.092 [0.009 – 0.182] |
| k₃GLU (min⁻¹) | 0.000 [0.000 – 0.005] | 0.000 [0.000 – 0.010] | 0.000 [0.000 – 0.004] | 0.001 [0.000 – 0.016] |
| k_mitoGLU (min⁻¹) | — | — | — | — |
| ***Derived Parameters*** | | | | |
| VTGLN (mL/g) | 0.144 [0.117 – 0.352] | 0.243 [0.171 – n/a] | 0.524 [0.367 – n/a] | 0.321 [0.238 – n/a] |
| VTGLU (mL/g) | 0.367 [0.221 – 0.445] | 0.210 [0.146 – n/a] | 0.360 [0.263 – n/a] | 0.171 [0.112 – n/a] |
| FluxGLS (mL/min/g) | 0.023 [0.018 – 0.027] | 0.015 [0.010 – n/a] | 0.037 [0.021 – n/a] | 0.018 [0.010 – n/a] |
| *Abbreviations: TNBC = triple-negative breast cancer (HCC1806); ER+ = estrogen receptor-positive (MCF-7); CI = confidence interval; n/a = CI upper bound not available from simulation output. k₃GLU values near zero reflect negligible direct glutamate-to-macromolecule incorporation.* | | | | |

| **Supplementary Table 3.** Bias and precision of kinetic parameter estimates* | | | | | | | | | |
| --- | --- | --- | --- | --- | --- | --- | --- | --- | --- |
| **single glutamate pool model** | | | | | **dual glutamate pool model** | | | | |
| **Bias (%)** | HCC1806 Vehicle | HCC1806 Treated | MCF-7 Vehicle | MCF-7 Treated | **Bias (%)** | HCC1806 Vehicle | HCC1806 Treated | MCF-7 Vehicle | MCF-7 Treated |
| K_1GLN_ | 5.5 | 12.0 | 2.6 | 6.4 | K_1GLN_ | -3.8 | 3.1 | -0.8 | -1.5 |
| K_1_/k_2GLN_ | 1.5 | -1.0 | -5.1 | -3.5 | K_1_/k_2GLN_ | 25.8 | -0.8 | 74.0 | 61.1 |
| K_1GLU_ | 24.4 | 45.4 | 127.0 | 90.0 | K_1GLU_ | 14.2 | 81.4 | -54.6 | -46.9 |
| K_1_/k_2GLU_ | 63.3 | -14.4 | -29.0 | 0.9 | K_1_/k_2GLU_ | 0.1 | 6.8 | -63.0 | -53.4 |
| $K_{1CO_{2}}$ | 21.5 | 1.2 | -17.2 | 6.9 | $K_{1CO_{2}}$ | 29.3 | -14.7 | -55.2 | -5.7 |
| $K_{1}/k_{2CO_{2}}$ | -4.9 | 0.4 | -27.1 | 6.1 | $K_{1}/k_{2CO_{2}}$ | 4.3 | 7.5 | 47.7 | 10.3 |
| k_3GLN_ | -4.3 | -4.8 | 12.7 | -2.6 | k_3GLN_ | 1.8 | 4.6 | -5.1 | -2.8 |
| k_GLS_ | -13.6 | -29.3 | -37.5 | -23.2 | k_GLS_ | -3.8 | -29.4 | 41.3 | 37.7 |
| k_TCA_ | 279.3 | 30.8 | 196.9 | 25.4 | k_TCA_ |  |  |  |  |
| k_GS_ | -16.5 | -50.7 | -39.5 | -33.7 | k_GS_ | -27.5 | -36.1 | -26.9 | -10.6 |
| k_3GLU_ | 14132.0 | 10372.0 | 17967.0 | 86.0 | k_3GLU_ | -55.9 | 65.1 | -5.1 | 971.0 |
| k_mito_ | - | - | - | - | k_mito_ | 1.8 | -44.6 | -37.7 | -47.2 |
| $V_{TGLN}$ | 13.2 | 19.2 | 36.6 | 15.4 | $V_{TGLN}$ | -3.7 | -7.0 | 0.9 | -5.1 |
| $V_{TGLU}$ | 8.2 | 31.3 | 39.6 | 27.7 | $V_{TGLU}$ | -1.6 | 6.6 | 19.9 | 4.9 |
| $V_{TGLU,mito}$ | - | - | - | - | $V_{TGLU,mito}$ | -7.9 | 13.9 | 125.7 | 135.4 |
| $V_{TGLU,cyto}$ | - | - | - | - | $V_{TGLU,cyto}$ | 23.0 | -0.4 | -21.6 | -36.7 |
| ${Flux}_{GLS}$ | -6.3 | -18.0 | -17.7 | -13.8 | ${Flux}_{GLS}$ | -7.2 | -33.8 | 42.8 | 30.2 |
| **CV (%)** | HCC1806 Vehicle | HCC1806 Treated | MCF-7 Vehicle | MCF-7 Treated | **CV (%)** | HCC1806 Vehicle | HCC1806 Treated | MCF-7 Vehicle | MCF-7 Treated |
| K_1GLN_ | 12.3 | 13.0 | 12.8 | 12.9 | K_1GLN_ | 9.1 | 16.0 | 11.4 | 13.4 |
| K_1_/k_2GLN_ | 35.3 | 30.4 | 27.4 | 29.4 | K_1_/k_2GLN_ | 38.6 | 30.2 | 50.7 | 64.0 |
| K_1GLU_ | 76.9 | 127.8 | 160.1 | 251.0 | K_1GLU_ | 46.8 | 192.5 | 31.0 | 48.7 |
| K_1_/k_2GLU_ | 67.0 | 47.2 | 37.6 | 57.0 | K_1_/k_2GLU_ | 94.2 | 89.5 | 35.7 | 46.0 |
| $K_{1CO_{2}}$ | 73.4 | 49.5 | 82.1 | 73.3 | $K_{1CO_{2}}$ | 95.6 | 25.4 | 61.8 | 53.0 |
| $K_{1}/k_{2CO_{2}}$ | 21.6 | 20.6 | 49.6 | 35.8 | $K_{1}/k_{2CO_{2}}$ | 16.6 | 23.6 | 134.4 | 28.3 |
| k_3GLN_ | 9.2 | 9.1 | 21.0 | 6.5 | k_3GLN_ | 5.5 | 11.6 | 12.0 | 4.6 |
| k_GLS_ | 13.2 | 25.0 | 23.4 | 20.2 | k_GLS_ | 10.5 | 17.9 | 54.7 | 55.3 |
| k_TCA_ | 416.8 | 123.2 | 396.3 | 131.8 | k_TCA_ | - | - | - | - |
| k_GS_ | 20.5 | 42.0 | 23.0 | 53.1 | k_GS_ | 32.1 | 29.4 | 27.9 | 44.9 |
| k_3GLU_ | 14806.0 | 4342.5 | 3747.7 | 116.8 | k_3GLU_ | 65.6 | 216.4 | 111.2 | 1211.2 |
| k_mito_ | - | - | - | - | k_mito_ | 1.9 | 28.1 | 38.3 | 27.2 |
| $V_{TGLN}$ | 13.3 | 21.2 | 27.8 | 23.0 | $V_{TGLN}$ | 10.6 | 20.0 | 10.2 | 11.2 |
| $V_{TGLU,mito}$ | - | - | - | - | $V_{TGLU,mito}$ | 13.8 | 34.0 | 84.6 | 93.4 |
| $V_{TGLU,cyto}$ | - | - | - | - | $V_{TGLU,cyto}$ | 43.7 | 40.1 | 27.2 | 28.4 |
| $V_{TGLU}$ | 12.1 | 25.5 | 21.7 | 35.1 | $V_{TGLU}$ | 15.7 | 22.6 | 12.8 | 16.7 |
| ${Flux}_{GLS}$ | 10.1 | 15.4 | 15.9 | 15.1 | ${Flux}_{GLS}$ | 14.1 | 24.5 | 54.8 | 53.2 |

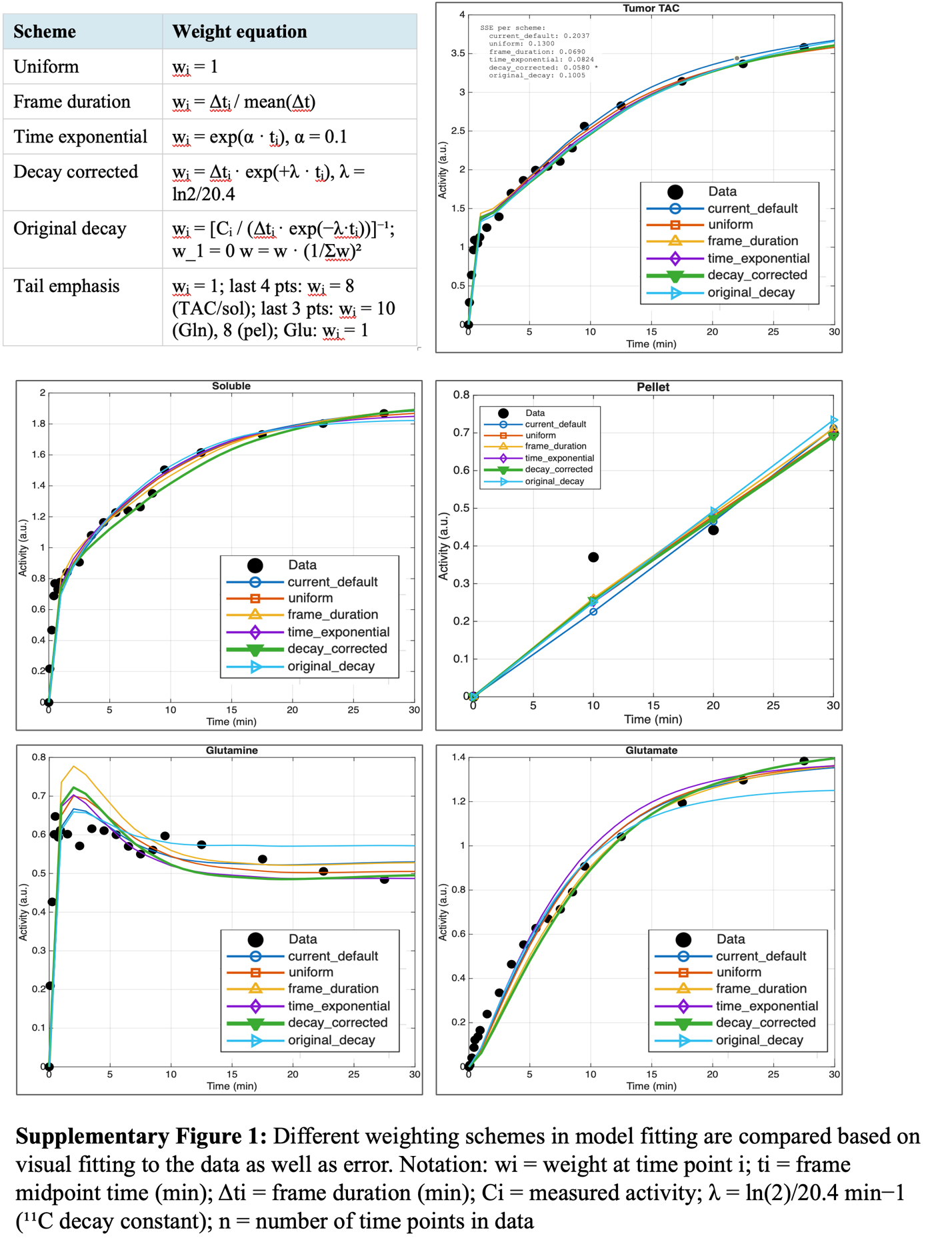

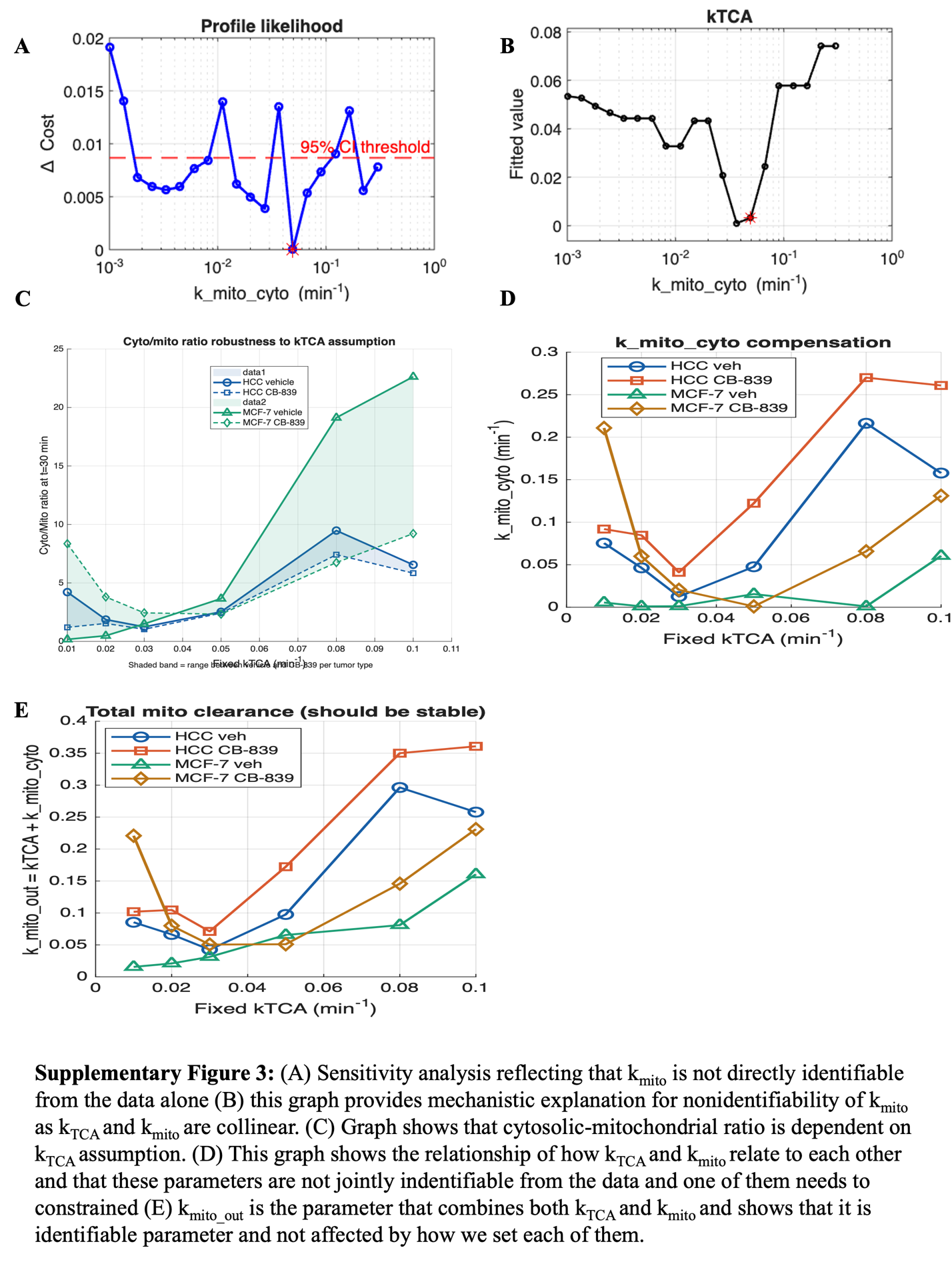

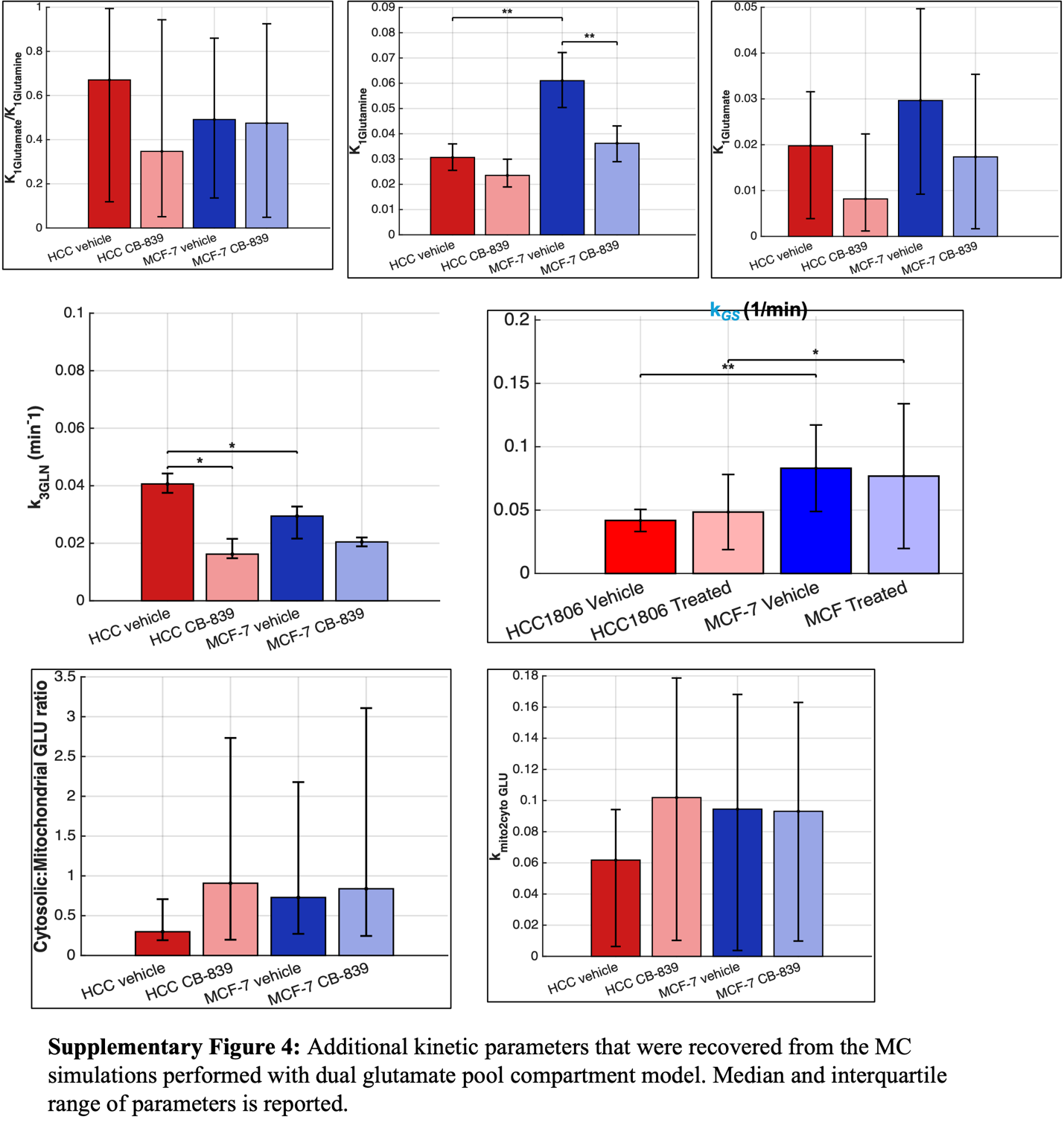
